# Multiplexed analysis of multicomponent biomolecular condensates without any tag

**DOI:** 10.1101/2025.10.10.681607

**Authors:** Gyula Pálfy, Johannes Schmoll, Maria E. Pérez, Fred F. Damberger, Yaning Han, Leonidas Emmanouilidis, Frédéric H-.T. Allain, Mihajlo Novakovic

## Abstract

Biomolecular condensates, formed in the process of liquid-liquid phase separation (LLPS), play key roles in RNA metabolism and cellular organization. These dynamic assemblies often contain several components such as proteins and RNAs. Experimental methods to study biological condensates commonly involve fluorophore-labelling of the biomolecules, which may affect phase separation behavior and in addition cannot access dilute-phase components due to low dye concentrations. Here we introduce a non-invasive, multiplexed NMR approach that enables selective observation of multiple components in hydrogel-stabilized biphasic condensates without using tags. The introduced multiplexing filter combines enhanced spin-spin cross-relaxation in the condensed phase with diffusion and isotope filters to resolve signals from distinct protein pools from both dilute and condensed phases. We demonstrate the robust performance of this 1D ^1^H NMR experiment using biphasic samples containing condensates of FUS N-terminal domain, FUS full-length, and the multicomponent condensate formed by two intrinsically disordered regions of hnRNPC1 protein with highly overlapping resonances. Integrating the multiplexing filter into multidimensional experiments enables site-specific structural and dynamic insights from diverse protein populations, extending NMR applications in LLPS.

## MAIN TEXT

Cells rely on the formation of biomolecular condensates to organize and regulate a crowded cellular milieu.^1,2^ Involved in RNA metabolism, biomolecular condensates are often enriched in proteins that contain RNA-binding domains (RBDs) and LLPS-prone intrinsically disordered regions (IDRs).^3–6^ Fluorescence microscopy has emerged as a powerful method of choice to study condensate morphology and dynamics both in cells and in vitro. However recent reports suggest that fluorescent tags can affect the LLPS behavior of biomolecules and thus bias interpretation,^7– 10^ especially in multicomponent systems where multiple spectroscopically distinct tags need to be used simultaneously for multicolor analysis of individual components. Furthermore, the dilute phase, the largest part of the overall volume, remains unobserved due to low dye concentration.

Solution NMR spectroscopy offers a tag-free methodology to probe structure, dynamics and interactions in complex environments. However, most NMR studies of biological condensates have focused solely on the macroscopic condensed phase, obtained by centrifugation of biphasic mixtures.^11–14^ Although providing important structural insights, this lacks the droplet morphology observed in vivo and does not capture the dynamic equilibrium between coexisting phases at their interface.^15,16^ As this interface can play a crucial role in condensate aging,^17–19^ associated with disease progression, it is important to study biomolecular condensates non-invasively in their native biphasic environment. Cytoskeleton-mimicking agarose hydrogels can effectively stabilize liquid droplets^16,20^ limiting their size to near-physiological ranges and allowing prolonged spectroscopic analysis without sedimentation. In addition, preparation of biphasic samples requires much smaller protein quantities compared to macroscopic condensed phase samples. However, in biphasic samples, the spectral overlap of ^1^H signals stemming from the two phases and different protein constituents complicates NMR analysis. Here we introduce a universal pulse sequence block with multiplexing capabilities that can select different components of biomolecular condensates and can be combined with most of the multidimensional NMR experiments used to probe structure and dynamics. Utilizing the very different cross-relaxation rates of the two phases, their diffusion contrast as well as isotope labeling, our multiplexing filter can resolve the spectra of at least four different molecular states in a biphasic multicomponent system, providing more complete insights into multicomponent condensates.

Facilitating phase separation, IDRs play a crucial role in biomolecular condensation and are often part of the proteins found in membraneless organelles.^3–5^ IDRs remain disordered in the condensed phase with side chains being locally highly dynamic resulting in comparable linewidths of their signals stemming from dilute and condensed phases. Additionally, their spectra are often highly overlapped.^12^ Given large differences in diffusion coefficient between the two phases (2-3 orders of magnitude),^12,16,21^ a diffusion filter can be very effective for selecting the signals originating from the slow diffusing molecules in the condensed phase,^20^ but an alternative filter for observing only the dilute phase remains challenging. So-called relaxation filters (i.e. T_2_ filter)^22^ to detect solely the component with slower relaxation are not very efficient at suppressing the condensed phase signals because their proton relaxation rates are similar to those of the dilute phase (Supporting Note 1 and Figure S1A). However, the cross-relaxation between the proton spins becomes highly efficient in the condensed phase because of the significantly increased intermolecular contacts and compaction in the dense phase, increasing the effective proton density (Figure 1A).^12,21,23–26^ In addition, the slower global rotational correlation time due to higher viscosity in the condensed phase also boosts the efficiency of cross-relaxation, the ensuing NOE effect^27^ and spin diffusion. This leads to very fast magnetization transfer (MT)^28,29^ in the condensate, allowing one to suppress the spins by irradiation of their spectroscopically-distinct, cross-relaxing neighbors while preserving most of the resonances of the dilute phase. For example, saturation of all amide protons is much more efficiently transferred to the aliphatic protons within the molecules in the condensed phase, compared to the dilute phase, reducing their intensity and providing an MT contrast^30,31^ between the phases (Supporting Note 1 and Figure 1B).

**Figure 1.**
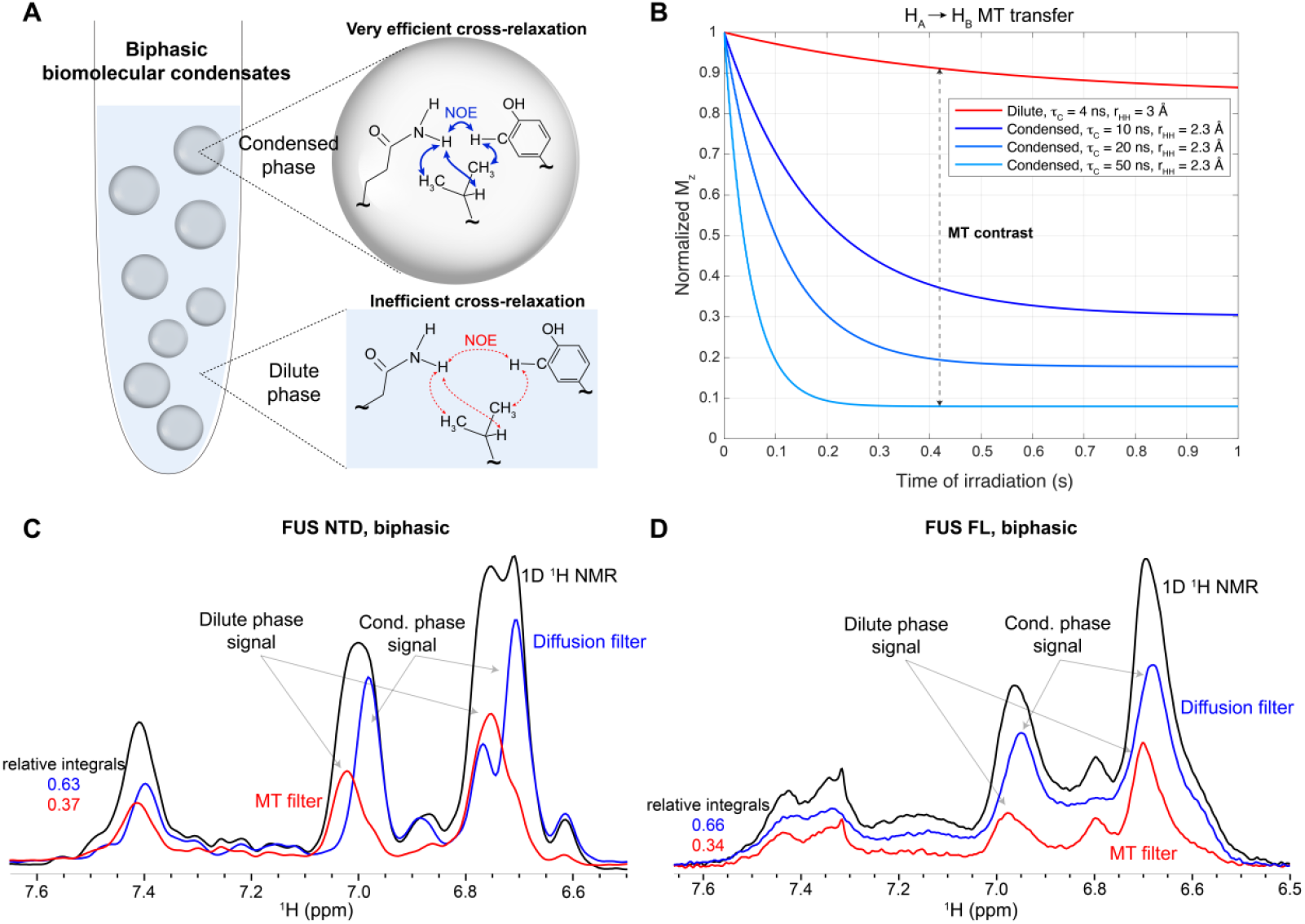
(A) Biphasic biomolecular condensates can be stabilized using 0.5% agarose hydrogel. Due to enhanced intra- and intermolecular contacts, slower molecular tumbling and compaction, cross-relaxation and the ensuing NOE effect is more efficient in the condensed compared to dilute phase. (B) Simulations performed using the Bloch-McConell-Solomon equations incorporating a continuous saturation of one proton pool (i.e. amide) and evaluating the ensuing steady state magnetization transfer to the other proton pool (i.e. aliphatic) through cross-relaxation. For simplicity, to depict the higher effective proton density and calculate generic cross-relaxation rates, we assumed several combinations of global τ_C_ and average interproton distance r_HH_ for medium-sized IDRs in dilute and condensed phase. Increase of τ_C_ and decrease in r_HH_ illustrate the effect of enhanced interactions upon condensation. (C,D) The efficiency of the so-called MT filter in selecting the dilute phase signal in FUS NTD and FUS FL sample. Condensed phase acquired using diffusion-filter is shown for completeness. The relative integrals of signals in condensed and dilute phase match well with corresponding protein populations validated using separate experiments. In the MT filter, aliphatic peaks at 0-1 ppm are irradiated with a nutation field of 1000 Hz.

The so-called MT filter has been tested on the biphasic sample of FUS NTD domain, comprising the low complexity QGSY-rich segment and the first arginine-glycine repeat (RGG1), which phase separates at micromolar concentrations and drives phase separation of the FUS protein.^32^ Despite largely overlapping signals in the sidechain aromatic region, the MT and diffusion filters can very efficiently resolve the FUS NTD signal stemming from the dilute and condensed phases (Figure 1C). As illustrated in Figure S1B, irradiation of aliphatic protons upfield from water selectively saturated 80% of the condensed phase signal while reducing only around 13% of the signal due to the saturation transfer in the dilute phase, similar to the predictions in Figure 1B. We further tested the performance of the MT filter on a FUS full-length (FUS FL) biphasic sample (Figure 1D). Here, the sum of the extracted dilute and condensed phase signals perfectly coincides with the total signal obtained without using filters (Figure S1C). Relative peak integrals of FUS NTD and FL in dilute and condensed phases also agree with protein populations determined by our REDIFINE approach (Figure S1D).^16^

Most biologically relevant condensates contain multiple components. This increases sample complexity as the coexisting dilute and condensed phases contain different pools of the protein or RNA molecules. To deconvolve different constituents, we combined diffusion and MT filters together with an isotope filter/edit block^33^ within a single pulse sequence (Scheme 1A). Utilizing differentially ^15^N- and ^13^C-labeled compounds, this multiplexing filter can be used in a 1D ^1^H experiment which allows the selection of the desired component of a biphasic sample by combining the different filters (Scheme 1B). The pulse sequence is described in detail in Supporting Note 2 and Figure S2.

**Scheme 1.**
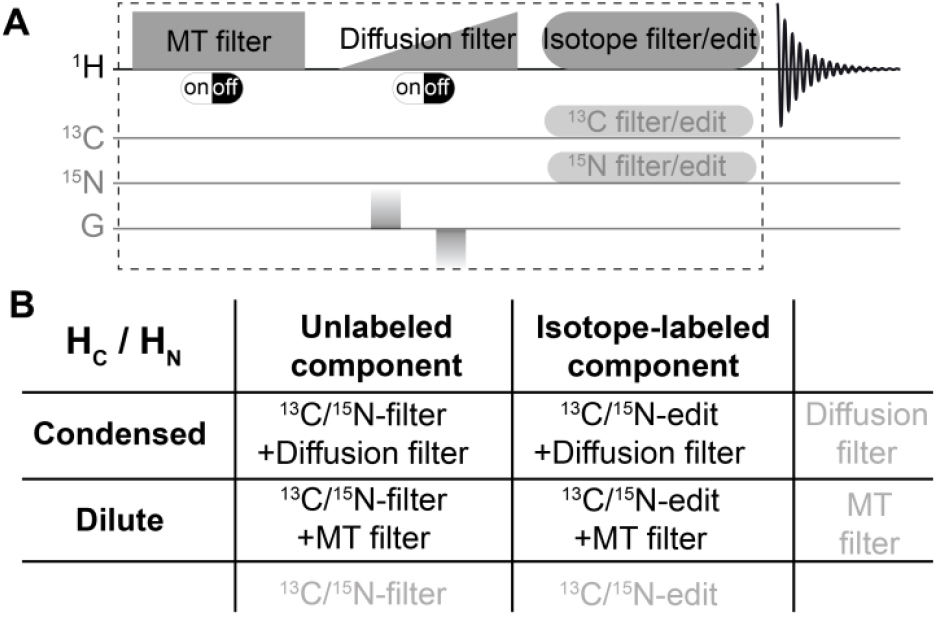
(A) Illustration of the multiplexing filter designed to resolve different components of complex biomolecular condensates. MT and diffusion filters can be switched on or off by turning on/off saturation or strong gradient, respectively. (B) Four different condensate constituents with overlapping signals can be selected using the appropriate combination of filters.

To test the filtering capability of this experiment on multiple-component condensates we mixed two intrinsically disordered domains of heterogeneous nuclear ribonucleoprotein C1 (hnRNPC1), namely IDR1 and IDR2 (Supporting Note 3 and Figure S3A). These constructs readily phase separate reaching the highest turbidity at a molar ratio of 2:1 (Figure S3B,C). Droplets were stabilized by preparation in 0.5% agarose. Two protein components (^15^N-labeled IDR1 and ^13^C-labeled IDR2) present in each phase yield four distinct protein states that can now be individually studied by NMR. Using our multiplexing filter, we were able to resolve the signals from each component even though their spectra fully overlapped (Figure 2A,B). Reassuringly, the sum of the signals stemming from aliphatic protons from dilute and condensed phases for both IDR1 and IDR2 almost perfectly match the spectrum acquired using only isotope filter/edit experiment.

**Figure 2.**
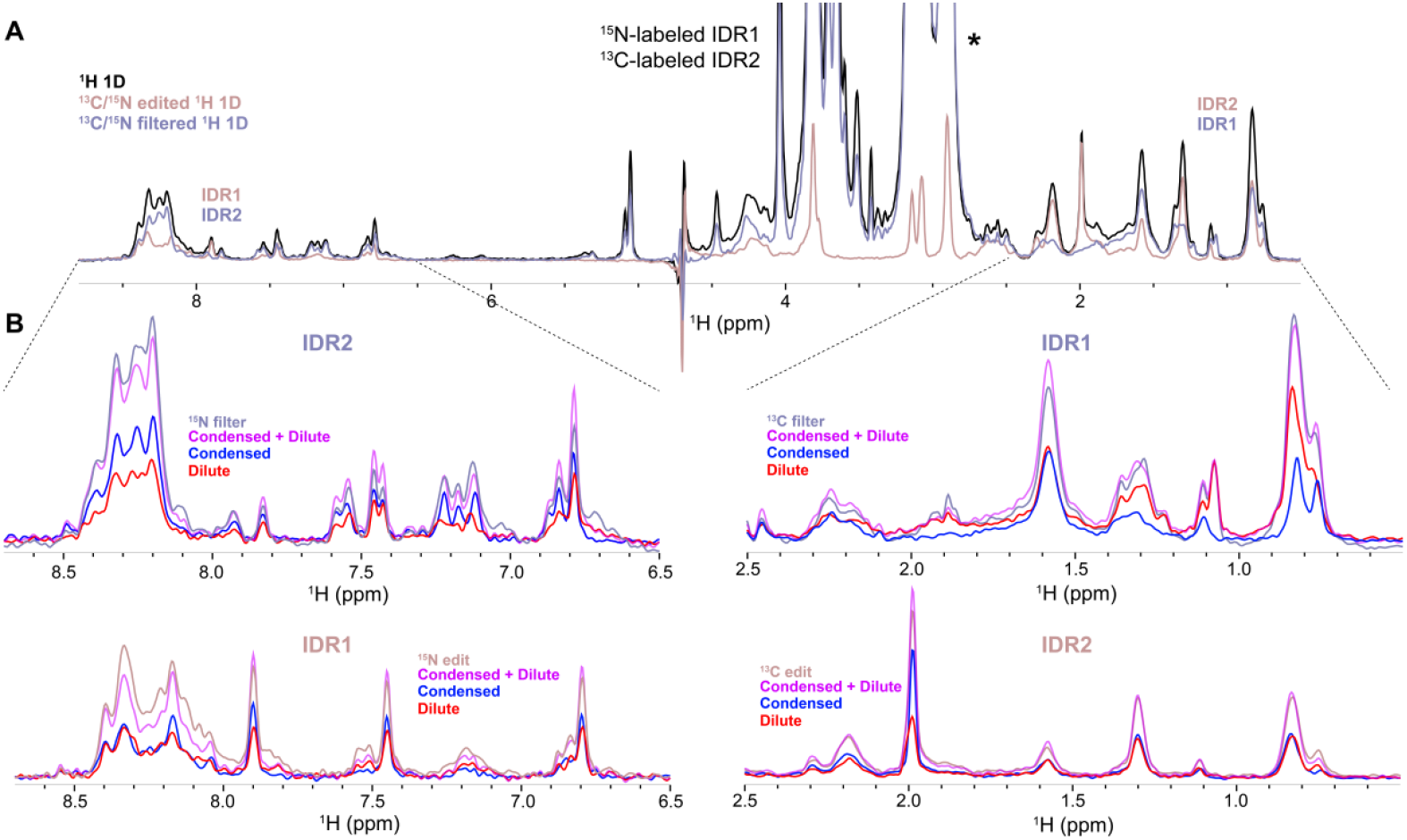
(A) Using an isotope-filter/edit block one can select different proteins according to their isotope enrichment. (B) Combined with isotope-filtering/editing, diffusion and MT filters can further resolve overlapping signals stemming from condensed and dilute phases. For amide detection, aliphatic proton peaks are saturated and vice versa. The efficiency of selecting the full signal from both phases is highly efficient for aliphatic signals. For amide detection, the presence of chemical exchange can affect discrimination of both phases, obtained with diffusion and MT filter, leading to an underestimation of the respective populations. Signals labeled with * are stemming from HEPES buffer.

Subtle chemical shift differences between the two phases are noticeable in the aliphatic region of IDR1 especially at 0.8 and 1.1 ppm demonstrating the high selectivity of the experiment.

While the signals in the aliphatic region perfectly sum up to the total signal, this is not the case in the amide region due to the presence of chemical exchange and more extensive broadening in the condensed phase. The backbone amides are typically broadened by the slower global tumbling in the condensed phase which, in addition to chemical exchange with water, can result in relaxation losses during the diffusion filter block. Furthermore, partial water saturation during the MT filtercan also be transferred to exchanging amides and lead to further underestimation of the dilute phase. Therefore, special considerations need to be taken when studying labile protons as discussed in Supporting Note 4.

Naturally, the multiplexing filter can be combined with other multidimensional NMR experiments to analyze the structure and dynamics of different protein pools. Unlike the diffusion filter, which introduces an additional relaxation pathway and is highly susceptible to relaxation and water exchange,^34^ the MT filter can be applied without relaxation losses (Figure S4A,B). This raised the idea to use the MT contrast to indirectly obtain the ^15^N-^1^H amide correlations stemming from the condensed phase by interleaving the irradiation between on- and off-resonance and performing difference spectroscopy. While the T_2_ contrast might be in principle similarly utilized for condensed phase selection, we observed similar linewidths for condensed and dilute phase amide proton signals in this particular system. Based on our simulations, this renders T_2_ filter experiment inefficient (Supporting Note 5 and Figure S5). We therefore integrated our MT-based filter within an ^15^N-^1^H HSQC experiment to acquire the dilute phase, and via MT difference spectroscopy the condensed phase spectrum. We recorded spectra at 28.2 T field strength (1.2 GHz ^1^H frequency) to exploit enhanced spectral resolution and a larger frequency separation (in Hz) of the cw saturation offset from the water resonance, thus minimizing spurious saturation (Supporting Note 1 and 4). Figure 3A,B shows ^15^N-^1^H correlations of IDR1 from the two phases at 298 K and 288 K respectively, resolved from highly overlapping spectra. Zoomed insets demonstrate excellent selectivity of the MT filter, especially at 288 K where the cross-relaxation becomes even more efficient due to slower dynamics. In both cases, much better SNR was obtained compared to the diffusion-filtered experiments (Figure S6). Given the results in Figure 2B, this approach is similarly applicable for observing the ^13^C-^1^H correlations in the condensed phase.

**Figure 3.**
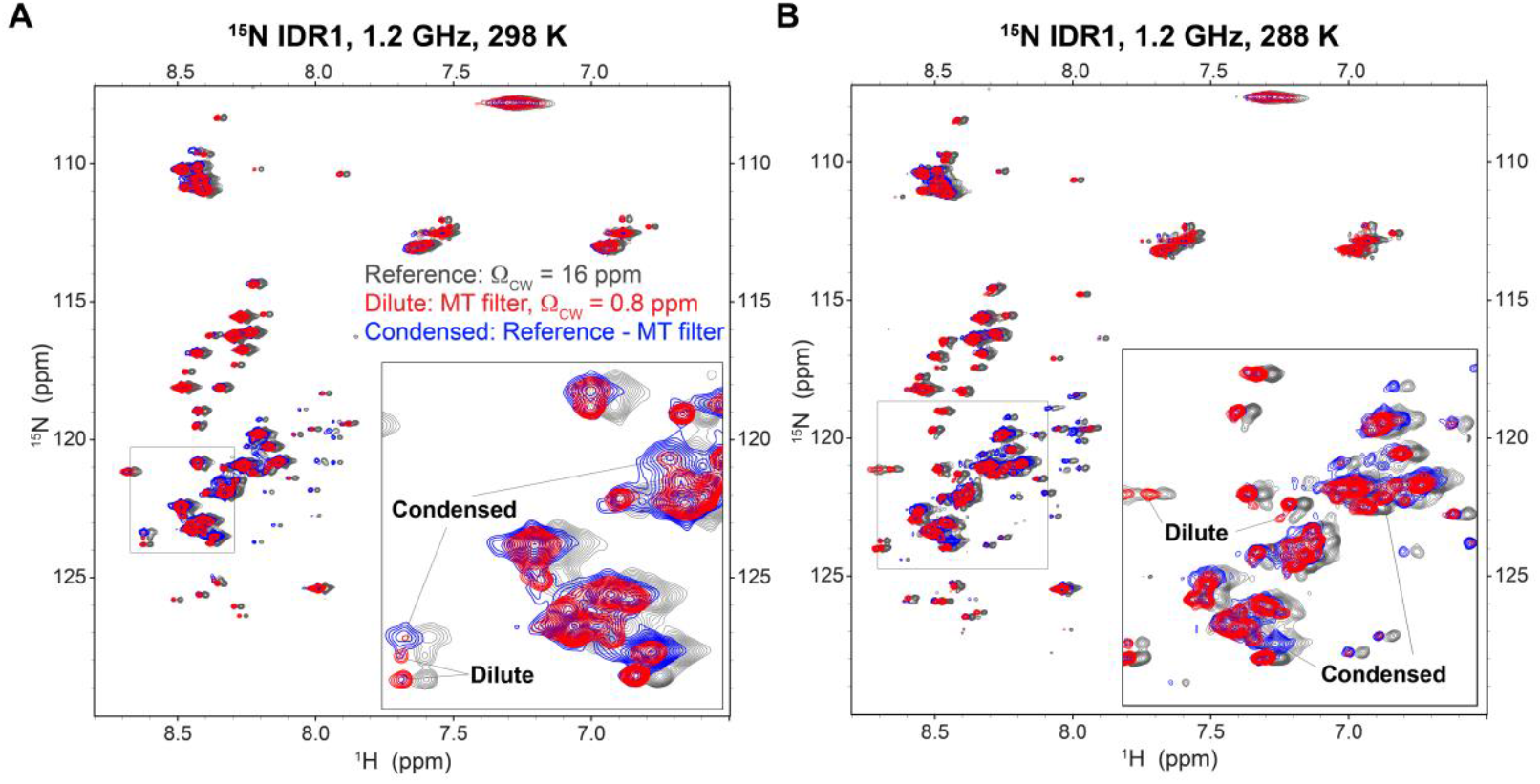
^15^N-^1^H correlations of IDR1 stemming from dilute and condensed phase acquired at (A) 298 K and (B) 288 K. Condensed phase spectra were obtained as a difference between MT-filtered HSQC and a reference spectrum where saturation is applied off resonance. The offset (16 ppm) was chosen to be downfield from amides at the same distance as the on-resonance saturation of aliphatic protons to correct for spurious saturation. Although heavily overlapping, zoomed insets highlight differences between dilute and condensed phase peaks. Reference spectra were arbitrarily shifted by −0.025 ppm in ^1^H dimension for clarity.

Our results demonstrate that the multiplexing filter can be used for “multicolor” analysis of protein-protein condensates resolving up to four components of a biphasic phase separated system. We envision that the multiplexing capabilities can be extended with yet another component which is a doubly ^15^N,^13^C-labeled molecule. In this case, for example, the aliphatic protons can be selected through multiple ^1^H-^15^N-^13^C-^1^H bond transfer, which might ultimately allow the analysis of up to six different molecular pools. Although not yet investigated, the same principle might be applicable to RNA molecules as these are very often an integral part of biomolecular condensates. The experiment is easy to set up, non-destructive and allows in-vitro analysis of biological condensates at near-physiological conditions. Discussed extensions of the current method and application in more complex environments are currently being investigated.

## ABBREVIATIONS

NMR: nuclear magnetic resonance
IDR: intrinsically disordered region
RBD: RNA-binding domain
LLPS: liquid-liquid phase separation
MT: magnetization transfer
HSQC: heteronuclear single quantum coherence

## ASSOCIATED CONTENT

Supporting Information (file type PDF)

## ACKNOWLEDGMENT

We would like to thank Dr. Alvar Gossert for incorporating excitation sculpting water suppression scheme within a time-shared double half filter, which we used as building block in this experiment.

