## Supporting Information for "Multiplexed analysis of multicomponent biomolecular condensates without any tag"

This PDF file includes:

Materials and Methods

Supporting Information Notes

Figures S1 to S6

Supporting Information References

### Materials and Methods

*Sample preparation*

FUS NTD low-complexity domain was expressed and purified as previously described.[1] The protein sample was prepared at the concentration of 200 μM for NMR analysis in 30 mM HEPES buffer with 200 mM KCl at pH 7.5 with a residual urea concentration of 0.6 M. The phase separation was induced by diluting 10x the 2 mM protein stock that was solubilized by 6 M urea. FUS FL protein was also expressed as previously described.[2] Similarly, 1.5 mM protein stock was diluted 10x to the final concentration of 150 μM for NMR analysis in 5 mM phosphate and 5 mM HEPES at pH 7, 1 mM TCEP, 0.6 M urea, and 15 mM NaCl. During the dilution step, agarose containing buffer at 55 ℃ was added to yield the final urea mass concentration of 0.5%. Agarose gelation occurs shortly after transferring to the NMR tube.[1]

IDR1 and IDR2 of human hnRNPC1 (Uniprot accession number P07910) were expressed in E.coli BL21 Star (DE3) or Rosetta (DE3) bacteria strains after transformation with the plasmids prepared by cloning the respective regions into pTEM vectors using $His_6$-GB1 tag (IDR1: 89-179 residues, IDR2: 207-293 residues of hnRNPC1). The cells were grown at 37 °C in M9 medium containing either $^{15}N$-$NH_4Cl$ and $^{12}C$-glucose (IDR1) or $^{14}N$-$NH_4Cl$ and $^{13}C$-glucose (IDR2) until $OD_{600}$ reached 0.6, then expression was induced by 0.1 mM IPTG for 5-6 h at 30 °C. Proteins were then purified by $Ni^{2+}$-affinity chromatography in 50 mM TRIS and 1 M NaCl (pH 8) buffer and eluted with gradually increased imidazole in 100-500 mM range, afterwards His-tag was removed by TEV protease overnight, while it was dialyzed against buffer containing 50 mM HEPES pH 6.5, 150 mM NaCl. The completely processed sample was applied to $Ni^{2+}$-affinity column again and the flow-through was collected, then concentrated to 3-4 mM and finally the stock solutions were stored at -20 °C. The NMR sample contained 400 μM $^{15}N$-labeled IDR1 and 200 μM $^{13}C$-labeled IDR2 in 50 mM NaCl in 50 mM HEPES, pH 6.0. During the mixing of the proteins agarose containing buffer was added at 42 ℃ resulting in a final concentration of 0.5%. Agarose gelation occurs shortly after transferring to the NMR tube.[1]

*NMR Spectroscopy*

All NMR experiments were recorded with a 3-mm NMR sample tube at 298 K or 288 K on 700 MHz Avance NEO spectrometers equipped with TCI cryo-probes with xyz-gradient system, 900 MHz Avance 3 HD equipped with TCI cryo-probe, and 1.2 GHz Avance NEO equipped with 3 mm TCI cryo probe. Experiments were run on TopSpin 3.6, 4.2 or 4.5, and all processed using TopSpin 4.5 software (Bruker).

The MT filter was performed using cw saturation with a duration of 0.5-5 s. Given the efficiency of cross-relaxation in the condensed phase, 0.5-1 s should suffice in most cases and should be used as a rule of thumb. When observing aliphatic protons, the cw irradiation offset was placed at around 8.5 ppm, and conversely for amide detection the offset was between 0 and 1.5 ppm. Irradiation at 0 ppm showed better performance. To ensure the proper bandwidth of saturation, the cw nutation field was varied from 500-1000 Hz. Although ca. 800 Hz was sufficient, we used 1000 Hz. To switch off the MT filter, far off-resonance saturation at -100000 Hz (-143 ppm) was used. For the reference scan used in $^{15}N$-$^{1}H$ HSQC difference spectroscopy at 1.2 GHz, cw saturation was applied at around 16 ppm, downfield from amides, to correct for spurious saturation effects arising from the on-resonance saturation of aliphatic protons.

The diffusion filter was performed using conventional stimulated echo with bipolar gradients.[3] To ensure that fast-diffusing signal stemming from the dilute phase was defocused, a diffusion gradient strength of 43 G/cm was used. In the case of FUS NTD, diffusion gradient time was 10 ms, with total diffusion time of 75 ms.[2] In the case of the multicomponent hnRNPC1 condensate we used 12 ms total gradient duration and 50-100 ms of diffusion delay. These numbers should be optimized for each specific sample using standard DOSY experiments. Smoothed rectangular-shaped gradients SMSQ10.100 were utilized.

A time-shared $^{15}N$/$^{13}C$ double half filter[4] was implemented which was combined with excitation sculpting[5] in the last part of the multiplexing block for water suppression. To calculate delays for scalar coupling evolution, 91 and 140 Hz were used as $J_{NH}$ and $J_{CH}$ coupling constants respectively. BUSS decoupling[6] was utilized on the $^{13}C$ channel while GARP4[7] was applied on $^{15}N$.

All 1D experiments were acquired using 32-128 scans and a $d_1$ recovery delay of 1.5 s. 2D $^{15}N$-$^{1}H$ HSQC correlation experiments were acquired using 32 scans (24 scans on 1.2 GHz), with 200 time increments (384 increments on 1.2 GHz) in the indirect dimension spanning 23 ppm (centered around 118.5 ppm) and $d_1$ of 1 s. For the MT filter, 1 s irradiation was used at the specified offsets. Diffusion-filtered $^{15}N$-$^{1}H$ HSQC was acquired using the same parameters as other $^{15}N$-$^{1}H$ HSQC experiments with the difference of using 128 scans to improve the SNR. On the 1.2 GHz, we used 24 scans for the Diffusion-filtered $^{15}N$-$^{1}H$ HSQC for direct comparison.

*Simulations*

Simulations on the MT and $T_2$-filter efficiency was performed using a home-written MATLAB codes that are available upon suitable request.

**Note 1: $T_2$ filter efficiency and magnetization transfer simulations**

Although there is a notion that protein signals significantly broaden in the highly dense phase in biomolecular condensates (as is the case with $^{15}N$ amide backbone resonances), the differences are less pronounced for $^{1}H$ signals, especially for intrinsically disordered regions of proteins. Preserving the fast local dynamics, side chains from IDRs usually remain highly dynamic even in the condensed phase. This leads to very subtle differences in both $T_1$[2] and $T_2$ relaxation parameters[8] between the two phases, rendering them practically indistinguishable based on relaxation parameters. Filtering out fast relaxing species by applying a spin echo is historically widely used, however the downside of this filter is that it monotonically suppresses also the desired slow relaxing signals. When relaxation constants are not significantly different as in the case of IDRs in condensates, this leads to a suppression of both species. This is simulated in Figure S1A. Two different scenarios of the combination of $T_2$ relaxation constants are assumed, realistic values of 50 and 33 ms (solid-colored lines) as well as a more favorable case of 80 and 20 ms (dashed-colored lines). Typically, one would choose 20, 40 or 70 ms echo delay (vertical dashed lines in Figure S1A). Clearly, in every case, a $T_2$ filter leads to significant reduction of signals from both phases and one can only achieve the near quantitative suppression of the condensed phase at 70 ms echo time. However, this would lead to an almost 60% reduction of the dilute phase, making the $T_2$-filter far from ideal for applications with biomolecular condensates.

On the other hand, Figure S1B illustrates the experimental performance of the MT filter applied to FUS NTD construct. Namely, by combining both MT and diffusion filter, one can assess the residual condensed phase that cannot be suppressed due to a steady state condition. We can see

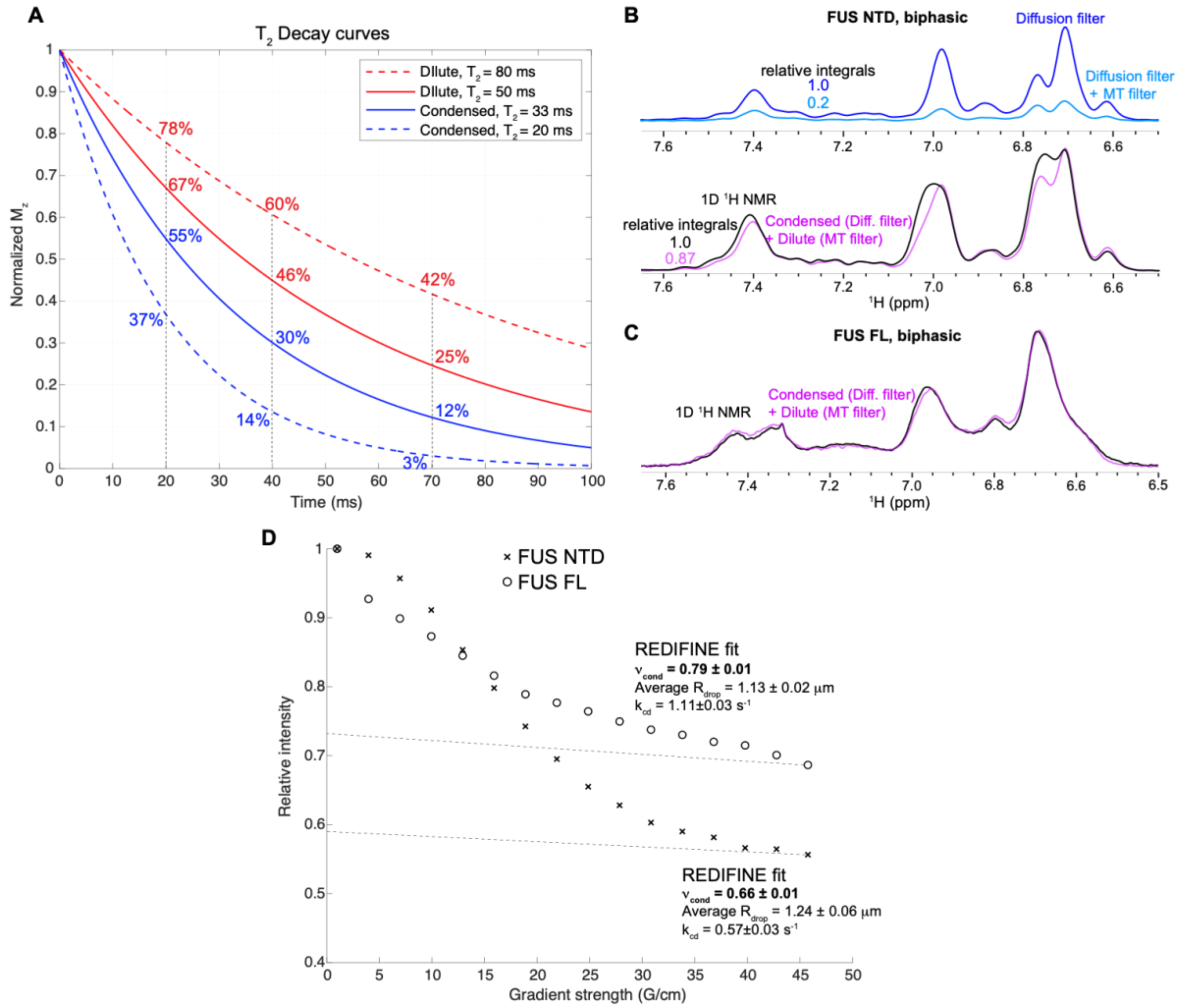


Figure S1. (A) Simulation of the filtering-efficiency of a $T_2$-filter experiment at various echo times and relaxation constants. Relaxation constants are realistically chosen according to the previously measured data and literature values. (B) Reduction of condensed phase signal upon application of an MT filter. As simulations in Figure 1B illustrated, MT can effectively suppress 80% of the condensed phase signal before reaching a steady state while reducing the dilute phase by ca. 15%. 5 s irradiation with a nutation field of 1000 Hz at 0.8 ppm (2720 Hz away from water at 700 MHz spectrometer) was used for the MT filter (C) The sum of the two phases is indistinguishable from the total $^1$H 1D signal (acquired using the same stimulated echo, but with diffusion gradients and MT filter off) for the FUS FL protein condensates. Here we used 3s irradiation with a nutation field of 1000 Hz, centered at -0.3 ppm (4500 Hz away from water at 900 MHz spectrometer). Signal decay in DOSY experiment (short 75 ms diffusion delay and 10 ms diffusion gradients) can be used to estimate the population of protein in condensed phase.[1] More accurately, REDIFINE method[2] accounting for the restricted diffusion inside the droplets and exchange was used to determine the exact populations of FUS NTD and FUS FL in the droplets.

here that MT filter effectively suppressed 80% of the condensed phase, akin to the simulation shown in Figure 1B that was performed using $t_C$ of 20 ns and $r_{HH}$ of 2.3 Å. Furthermore, from the difference between a total 1D $^1$H signal and the sum of the selected signal from the two phases (Figure S1B), we can see that the dilute phase is slightly underestimated by ca. 13% due to the much less efficient but still present cross-relaxation within the protein in dilute phase. This also matches with the prediction in Figure 1B using $\tau_C$ of 4 ns and $r_{HH}$ of 3 Å. Surprisingly, a similar comparison yields almost indistinguishable spectra for the FUS FL protein (Figure S1C), showing that MT and diffusion filter can in principle select the signal from the dilute and condensed phase without losses. To exclude relaxation losses, note that we refer to the total 1D $^1$H signal as the one acquired using the same length echo time, but with gradients and cw saturation being switched off. The better performance in the case of FUS FL protein might arise from the fact that MT contrast between dilute and condensed phase is more pronounced for FUS FL, containing also structured domains. Furthermore, application of cw at higher magnetic fields (900 MHz for FUS FL vs 700 MHz for FUS NTD) achieves similar saturation efficiency (as it is long enough to reach steady state) but is more selective due to the larger distance in Hz from water and other protein resonances. This effectively reduces spurious self-saturation of water and protein signal increasing the overall efficiency of the MT filter.

To check if the relative signal intensities shown in Figure 1C,D represent the relative populations of the protein in the condensed and dilute phase, we performed REDIFINE measurements.[2] While the data for FUS NTD match with the signal intensities determined from diffusion and MT filter experiments, the presence of faster exchange for FUS FL, mixing the components of the two phases, causes slight underestimation of condensed phase in Figure 1D. It is important to mention that fast chemical exchange between phases will intrinsically limit the selectivity of the MT filter, as the saturated protein in condensed phase can exchange to dilute phase and vice versa.

To simulate the differences in MT efficiency between the two phases we utilized a Bloch-McConnell-Solomon model[9] involving a two-spin system. This involved two protons $H_A$ and $H_B$ representing nearby aliphatic and aromatic/amide protons, connected via a generic cross-relaxation process, where one of them has been saturated using cw irradiation. The resulting equations can be written as:

$$\frac{dM_y^A}{dt} = w_{1A} M_z^A - R_2^A M_y^A$$

$$\frac{dM_z^A}{dt} = -w_{1A}M_y^A - (R_1^A + \sigma)M_z^A + \sigma M_z^B + R_1^A M_{eq}^A$$

$$\frac{dM_z^B}{dt} = -(R_1^B + \sigma)M_z^B + \sigma M_z^A + R_1^B M_{eq}^B$$

where $M_y^A$, $M_z^A$ and $M_z^B$ are the magnetization components of the proton spins along the specified axis of the Bloch sphere and $M_{eq}^A$ and $M_{eq}^B$ correspond to the equilibrium magnetizations of these reservoirs (for simplicity normalized to unity). Longitudinal and transverse relaxation rates were calculated as the inverse of the corresponding relaxation times.

The dipole-dipole cross-relaxation rate describing the magnetization transfer between two spins is given by

$$\sigma = \frac{1}{10}b^2(\mathcal{J}(0) - 6\mathcal{J}(2w^0))$$

where $\mathcal{J}(w) = \frac{\tau_c}{1+\omega^2\tau_c^2}$ is the spectral density function, and $b = -\frac{\mu_0}{4\pi}\frac{\hbar\gamma^2}{r_{HH}^2}$ is the dipole-dipole coupling constant. The strength of the saturation field applied along the x-axis is denoted as $w_{1A}$. For the simulation shown in Figure 1, we used the following parameters: $w_{1A} = 500\ Hz$, $T_1^A = 0.3\ s$, $T_1^B = 0.6\ s$, $T_2^A = 0.05\ s$, $r_{HH} = 3$Å or 2.3Å an arbitrarily chosen value assuming compaction upon condensation, $\tau_c = 4\ ns$ for dilute phase and $\tau_c = 10 - 50\ ns$ to effectively simulate the increase intra- and intermolecular contacts in the condensed phase, and $B_0 = 16.3\ T$ ($\omega H = 700\ MHz$).

**Note 2: Multiplexing filter pulse sequence – practical guide**

The pulse sequence for the multiplexing filter is shown in Figure S2. Briefly, it incorporates an MT filter (saturation to suppress condensed phase), diffusion filter (strong gradient to suppress dilute phase), and isotope filter/edit block to select isotope unlabeled or labeled components. All filters can be switched on and off depending on what is desired to be observed (Scheme 1 – main text). Solvent is suppressed with excitation sculpting during the last block. Importantly, the length of the entire multiplexing block was kept constant when acquiring different molecular pools, ensuring a constant total attenuation due to relaxation, assuming $T_2$ $^1H$ of all states are similar. This preserves the ratio of integrals of the different components in their individual spectra.

MT filter

If aliphatic signals upfield from water are desired, then the cw irradiation can be applied on amides downfield from water. As a rule of thumb, the irradiation can be centered between 8 and 9 ppm and the nutation field applied with $B_1$ field strength of 700-1000 Hz, for 0.5-1 s. To ensure the steady state, we applied the irradiation for 3-5 s in Figure 1, however this is usually not necessary. In Figure 2, we used 1 s irradiation. This achieves effective suppression of the condensed phase in the aliphatic region without pronounced self-saturation of the aliphatic peaks. Similarly, if aromatic and/or amide protons are desired, cw irradiation should be placed at 0 or 1 ppm which can effectively suppress the aliphatic protons. The MT filter selectivity benefits from higher fields due to reduced spurious water saturation (larger absolute frequency difference from water).

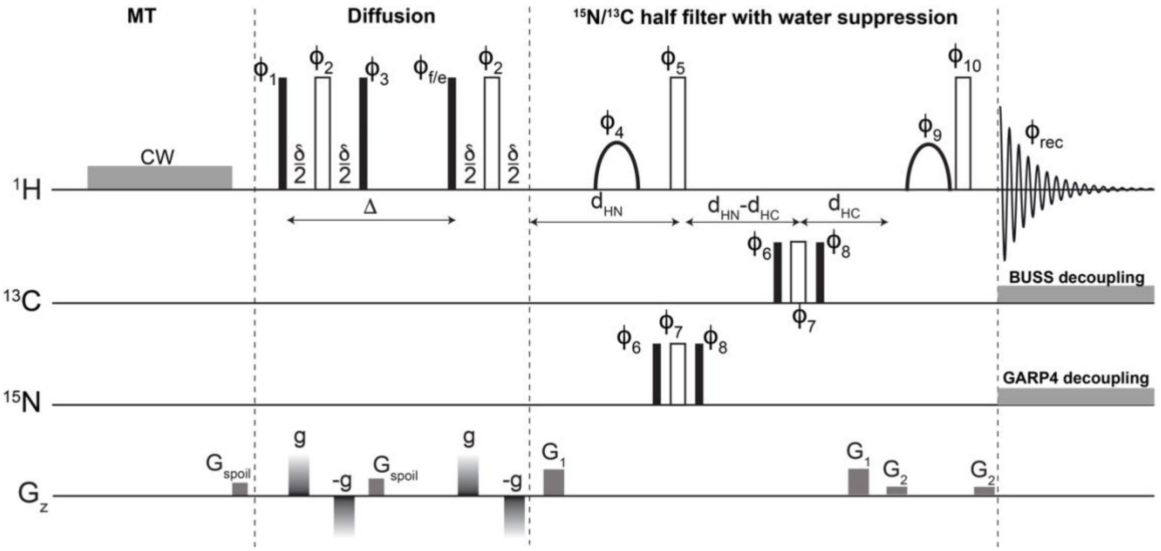


Figure S2. Pulse sequence for the multiplexing filter. Filled and outlined rectangles represent $\pi/2$ and $\pi$ pulses respectively (except the outlined pulses on the X channels which are defined below). $\Delta$ is the diffusion delay while $2\delta$ is the total duration of 4 gradients during the stimulated spin-echo. Delays are defined as $d_{HN}=1/2J_{NH}$ and $d_{HC}=1/2J_{CH}$. Outlined rectangles on $^{13}C$ and $^{15}N$ channels are defined as 2.66x the $\pi/2$ pulse as a part of a composite 180 pulse. The two shaped pulses are selective X shape Y ms $\pi$ pulses on the water resonance which in combination with the $G_1$ and $G_2$ gradients achieve excitation sculpting for solvent suppression. Phases are defined as: $\phi_1$=x, $\phi_2$=x, $\phi_3$=x, and depending on whether filter (f) or edit (e) flag is selected $\phi_f$=8(x), 8(y), 8(-x), 8(-y) or $\phi_e$=4(x), 4(-x), 4(y), 4(-y), 4(-x), 4(x), 4(-y), 4(y), $\phi_4$=x, y, $\phi_5$=-x, -y, $\phi_6$=x, $\phi_7$=y, $\phi_8$=4(x), 4(-x), $\phi_9$=x, x y, y, $\phi_{10}$=-x, -x, -y, -y, $\phi_{rec}$=2(x, -x, -x, x), 2(-y, y, y, -y), 2(-x. x, x, -x), 2(y, -y, -y, y). $G_{spoil}$ gradients serve to suppress residual transverse magnetization (1 ms each of 40% and -17.13% of the maximum amplitude, respectively), g is the variable gradient used in the stimulated echo of the diffusion filter (2% when off and 95% when on), while $G_1$ and $G_2$ are the parts of excitation sculpting scheme (1 ms at 31% and 11%). The half-filter with integrated excitation sculpting is based on a pulse sequence written by Alvar Gossert (ETH Zurich).

*Diffusion filter*

Parameters of the diffusion filter vary greatly with the protein length. As a rule of thumb, to filter out dilute phase signals, one can apply 50-75 ms diffusion delay and 10-12 ms gradient duration at >40 G/cm depending on the probe capabilities. To limit the effect of chemical exchange (dilute-condensed protein and proton exchange with bulk water), the diffusion delay should be as short as possible. When the $T_2$ relaxation of the protein of interest is fast, the standard 10-12 ms gradient duration dictating the minimum echo time can lead to significant loss of signal – in these cases shorter gradient duration should be chosen at the expense of longer diffusion time (relying on longer longitudinal relaxation). In other words, there is an interplay between these two parameters that need to be optimized depending on the system. Optimization of these parameters can be performed using a standard stimulated echo DOSY experiment.

*$^{15}N/^{13}C$ filter/edit block*

We used a time-shared double half filter[4] that serves as $^{15}N/^{13}C$ filter/edit block resulting in the cleanest selection according to the isotope-label while optimizing sensitivity. The delays are tuned according to protein one bond amide $J_{NH}$ and aliphatic $J_{CH}$ couplings. Composite pulses are used on the X channels. The half filter is combined with an excitation sculpting scheme for efficient water suppression. BUSS and GARP4 decoupling are utilized for $^{13}C$ and $^{15}N$ channels respectively.

**Note 3: hnRNPC1 IDRs readily phase separate**

hnRNPC1 is a crucial RNA-binding protein involved in various aspects of RNA metabolism. Besides the RRM and leucine zipper domains, this protein contains two disordered regions, IDR1 and IDR2 (Figure S3A). When IDR1 and IDR2 are mixed, they readily phase separate with the highest turbidity at the ratio of 2:1. As a model system, we prepared a biphasic sample of these condensates stabilized by 0.5% agarose gel using $^{15}N$-labeled IDR1 and $^{13}C$-labeled IDR2 (Figure S3B,C), both of which remain observable in $^{1}H$ NMR upon phase separation and undergo slow chemical exchange between phases.

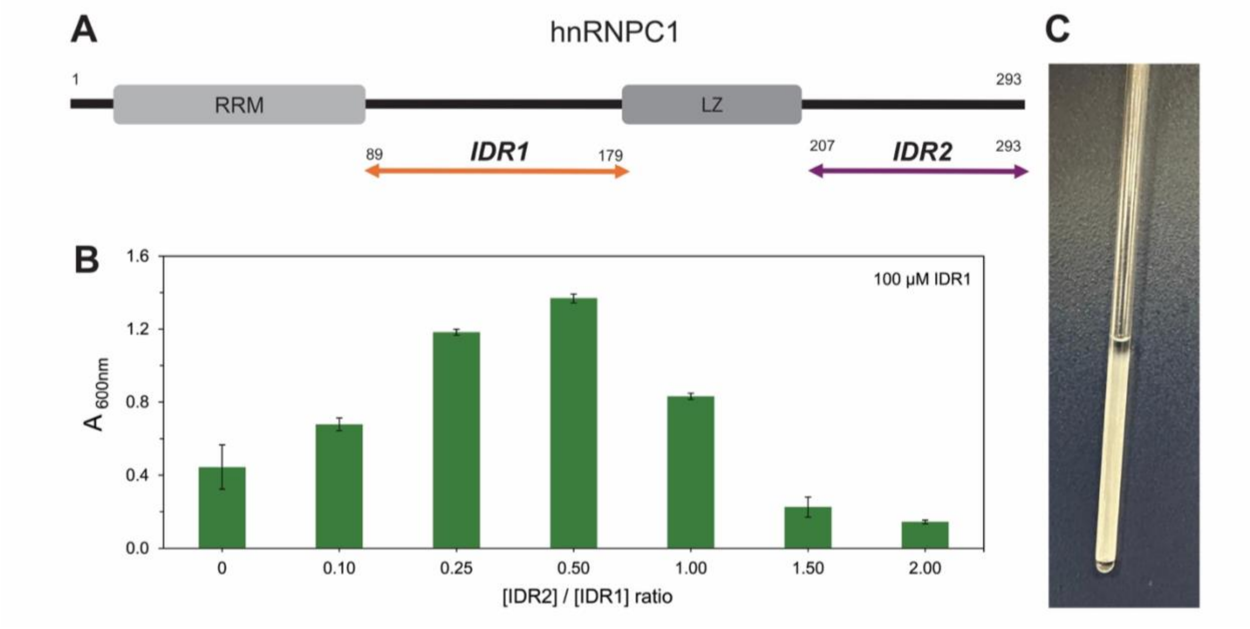


Figure S3. (A) Domain organization of the hnRNPC1 protein, illustrating the IDR1 and IDR2 domains used in this study. (RRM: RNA recognition motif, LZ: leucin-zipper domain, IDR1 and IDR2: disordered regions) (B) Turbidity assay of different IDR2/IDR1 molar ratio in the range of 0.0-2.0 using 100 μM IDR1 in 50 mM NaCl, 50 mM HEPES, pH 6.8. They undergo LLPS when mixed yielding the highest turbidity at the IDR1/IDR2 ratio of 2:1. (C) A biphasic NMR sample (400 μM $^{15}$N-IDR1, 200 μM $^{13}$C-IDR2 in 50 mM NaCl, 50 mM HEPES, pH 6.0) is shown which was prepared in 0.5% agarose, in which pH was lowered to decrease the water exchange to obtain sharper signals in the spectra.

**Note 4: Chemical exchange can complicate the diffusion and MT filter**

As already discussed in Note 2, chemical exchange between two phases and with bulk water can be detrimental for the efficiency of the diffusion filter.[10,11] This is why special care needs to be taken when working with amides. The potential problem can be visualized when comparing a diffusion-filtered $^{15}$N-$^{1}$H HSQC with the regular $^{15}$N-$^{1}$H HSQC. One can immediately see that exchanging amide signals decay somewhat faster in the diffusion DOSY experiment,[10,11] however the real problems can be pinpointed in a 2D spectrum (Figure S4A). Note that the diffusion-filtered HSQC suppresses also many peaks that can be associated with the condense phase and preserves only the signals with slowest $T_2$ relaxation and exchange. The combination of 50 ms diffusion delay and 12 ms gradient duration was used for the diffusion filter.

Similarly, chemical exchange with water can affect the MT filter when selecting amides. Namely, spurious water saturation can be transferred to exchanging amide protons underestimating the signals stemming from dilute phase. It is therefore beneficial to irradiate at an offset as far as possible from water to reduce this effect (Figure S4B). This can be appreciated by the increase of the dilute phase signal and the fact that the sum of dilute phase and condensed phase matches better

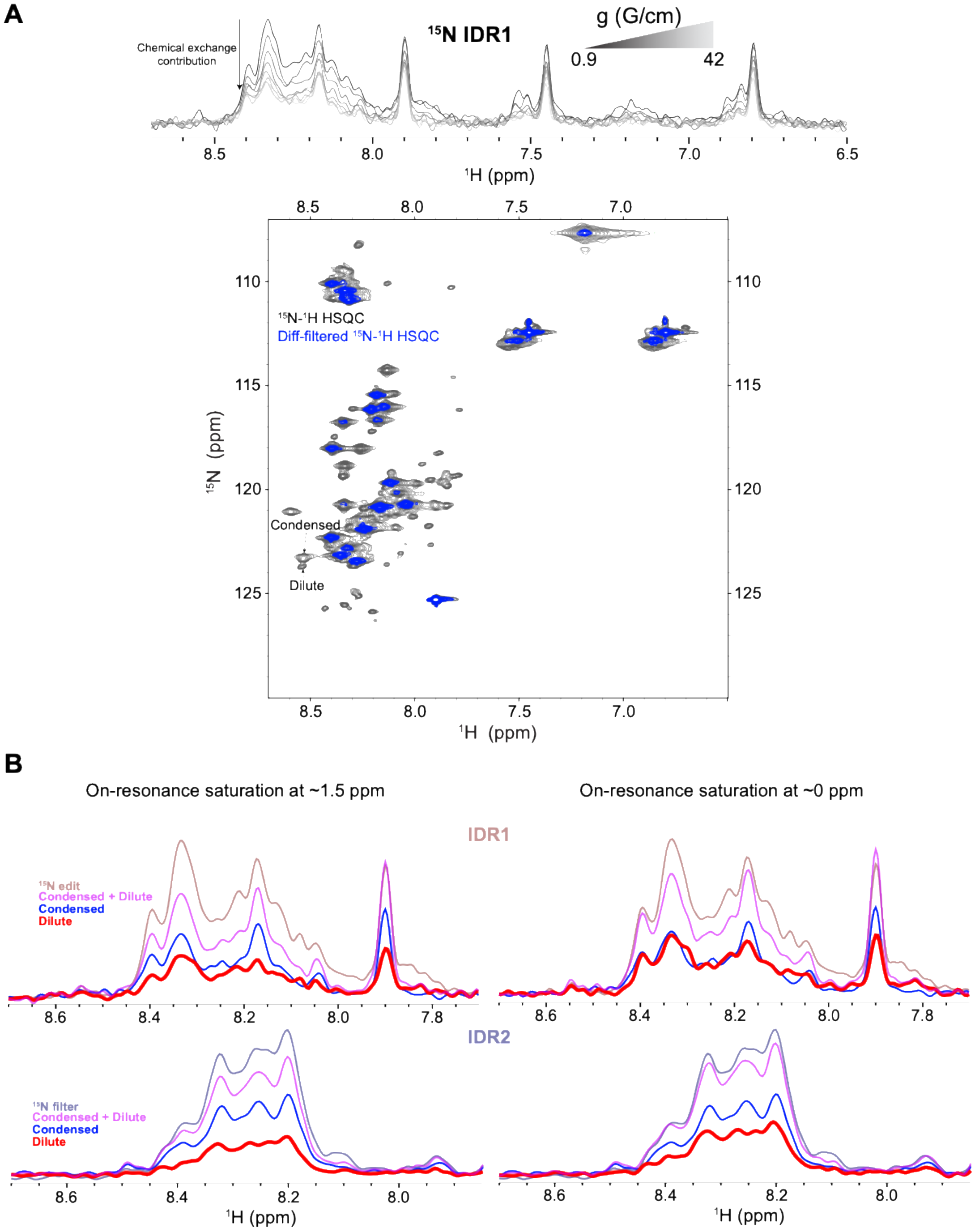


Figure S4. (A) $^{15}$N-edited DOSY decay of IDR1 as a part of a condensate with IDR2 and the corresponding diffusion-filtered $^{15}$N-$^{1}$H HSQC. Note that many signals are fully suppressed, even the ones that are known to stem from the protein in the condensed phase. (B) Shifting the irradiation from 1.5 to 0 ppm in the MT filter enhances the amides in the dilute phase due to the lesser extent of spurious water saturation.

with the total proton signal when irradiation is performed at 0 ppm instead of 1.5 ppm. This is why higher fields are especially beneficial for the MT filter as illustrated in Note 1 on the example with FUS FL. At higher magnetic fields, aliphatic protons resonate further from water, allowing one to irradiate far enough from water, while still effectively saturating the protein (1 ppm on 700 MHz is 2600 Hz from water, while 3300 Hz on 900 MHz and 4400 Hz on 1.2 GHz). In addition, it should also be noted that fast interphase exchange of biomolecules (mixing the content of the two phases) is an intrinsic limitation for the selectivity of both filters. However, unstructured proteins forming condensates generally undergo slow interphase exchange.[2]

**Note 5: $T_2$ filter applicability on amide protons of IDR1**

As pointed in the main text, a condensed phase signal indirectly extracted from the difference of an MT-filtered and a reference spectrum might provide improved sensitivity and fewer signals suppressed due to spurious water saturation compared to diffusion filtered HSQC. If cw irradiation is executed first on-resonance with protein and then interleaved by an “off” reference acquisition with a saturation pulse of the same $B_1$ intensity applied off resonance, one could acquire the condensed phase signal by calculating the difference spectrum of the on and off resonance saturation experiment. If amide protons have significantly different linewidths in the dilute and condensed phase, a $T_2$ contrast might be exploited in a similar manner. We identified in a $^{15}N$-$^{1}H$ HSQC spectrum of the $^{15}N$-labeled IDR1 in biphasic mixture with IDR2 two resolved peaks, stemming from the same residue from the two different phases (Figure S5A). From the extracted 1D slices containing these resonances (Figure S5A inset) we could calculate that the condensed phase peak is approximately 50% broader, approximately 30 Hz compared to 20 Hz in the dilute phase. Based on these linewidths, we used the corresponding $T_2$ relaxation constants to simulate the performance of the $T_2$ filter (Figure S5B). Note again that even with backbone amide protons there is no ideal spin-echo delay to effectively select the pure dilute phase component.

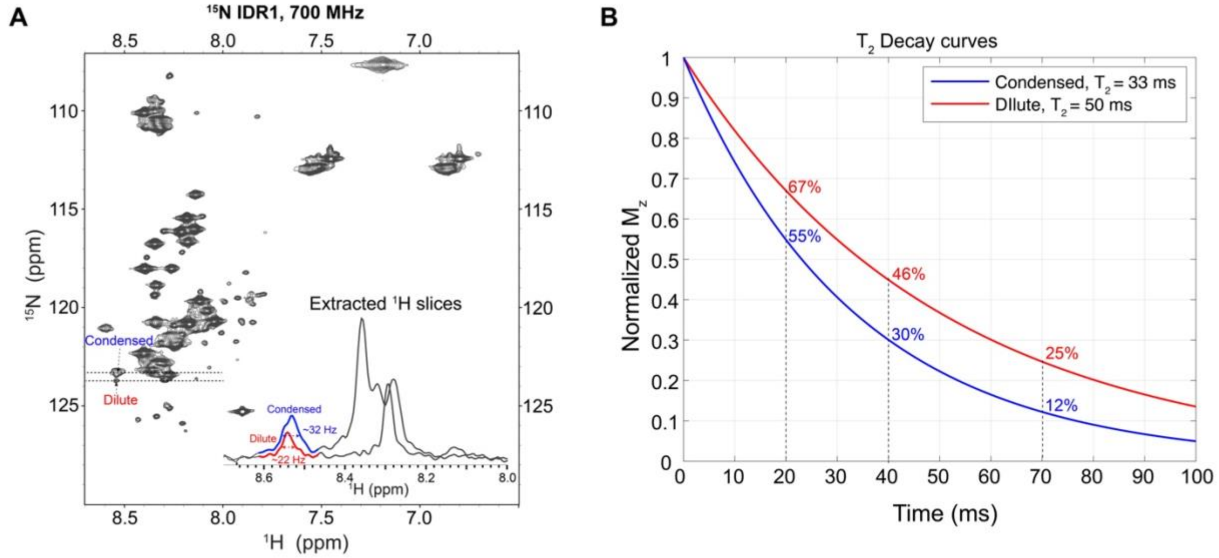


Figure S5. (A) $^{15}N$-$^{1}H$ HSQC of $^{15}N$-labeled IDR1 in the biphasic sample with IDR2. The dashed lines point to spectroscopically resolved resonances stemming from the dilute and condense phase signal of the same residue. 1D slices extracted from the specified $^{15}N$ chemical shifts (123.3 and 123.7 ppm), illustrating slightly different linewidths of amides in the two phases (~22 Hz vs. ~32 Hz), are shown in the inset. (B) Given the amide linewidths estimated in (A), we can simulate the $T_2$ decay showing that $T_2$ filter is not ideal choice to select the dilute phase of this particular sample.

**Note 6: The quality of the 2D amide condensed phase spectra**

Besides chemical exchange, fast transverse relaxation during stimulated echo period can further limit diffusion filter performance. This is especially pronounced in diffusion filtered heteronuclear correlation experiments where magnetization is subjected to additional coherence transfers. Figure S6A,B shows the comparison of diffusion-filtered $^{15}N$-$^{1}H$ HSQC spectra compared to the ones obtained using MT filter and difference spectroscopy at 298 K and 288 K. While the latter yielded the spectra of high quality at both temperatures, the diffusion-filtered spectra had very low SNR with the detrimental effect of fast $T_2$ relaxation especially pronounced at 288 K. On the contrary, slower dynamics at 288 K rendered cross-relaxation even more efficient, which further improves the performance of the MT filter illustrating that changing the temperature can further enhance the efficiency of the multiplexing filter.

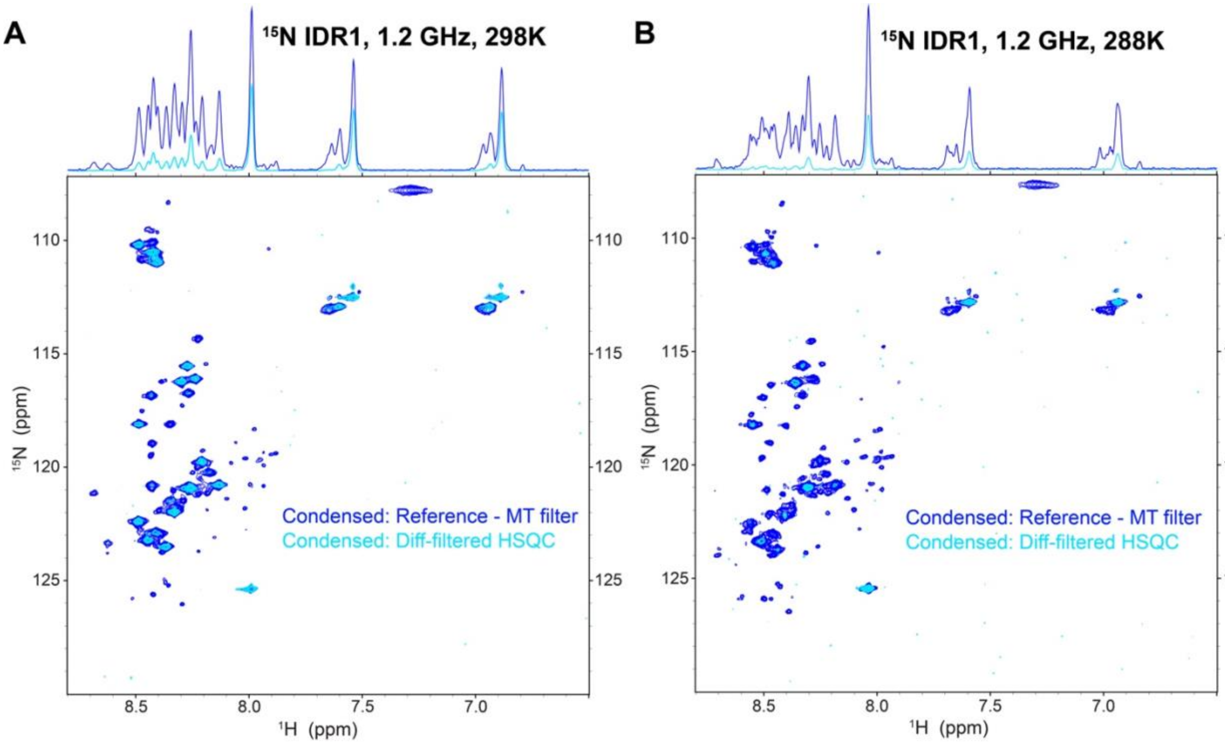


Figure S6. Comparison between the condensed phase $^{15}$N-$^{1}$H HSQC spectrum of IDR1, obtained by MT filtered difference spectroscopy and the diffusion filter, at (A) 298 K and (B) 288K. Although requiring 2-times more scans for acquisition, the condensed phase extracted from difference spectroscopy has far superior performance compared to the diffusion filter. This is especially pronounced at 288 K where faster $T_2$ relaxation due to slower tumbling proves detrimental for diffusion filter (due to the presence of relatively long echo). 1D spectra on the top represent $^{1}$H positive projections.
